# Small representative samples can capture global vascular plant diversity patterns

**DOI:** 10.64898/2026.07.08.737287

**Authors:** Ludwig Baldaszti, Peter W. Moonlight, Neil Brummitt, Samuel Pironon, Tiina Särkinen

## Abstract

Incomplete information on distributions for a high proportion of the world’s plant species together with biases in global biodiversity data mean that current estimates of plant diversity patterns are skewed. A key issue is that current predictions rely on a subset of species that is not representative of all plant species. Here we tested the feasibility of a representative sampling approach for mapping global vascular plant diversity at the finest scale where comprehensive data is available. Using the World Checklist of Vascular Plants as a reference, we generate random samples of species with increasing sample sizes from the global species pool. We compare the diversity patterns retrieved from the samples against the patterns of the reference dataset using spatially weighted correlation coefficients and four different diversity metrics. We find that at the botanical country scale, representative global maps of species and phylogenetic diversity can be created with small numbers of species (∼1% [0.2% and 0.4%, respectively]) at the botanical country scale. For effective growth form and family diversity sample sizes encompassing ∼20% [19.2% and 19.5%, respectively] of all species are needed. Random samples require markedly fewer species to reach high correlations than when restricting the pool of species to single plant families or genera. We show that when representative samples are used robust inferences of plant diversity patterns can be made from only a small proportion of species.

## Introduction

In the face of the accelerating biodiversity crisis, there arguably has never been a more important time to produce accurate maps of global biodiversity to facilitate conservation efforts. This is particularly true in light of the Kunming-Montreal Global Biodiversity Framework, adopted by 196 countries and aiming to accelerate and upscale the protection of biodiversity globally, including a target to protect 30% of the planet by 2030 (“30-by-30”) with a particular focus on the most biodiverse regions (CBD, 2022). Despite general agreement at broad global scales (Brummitt et al., 2021), however, identification of areas with high plant diversity at fine spatial scales suitable for implementing conservation policies and comparing their effect is still subject to debate.

In taxonomic groups such as vertebrates, expert-verified IUCN range maps, originally designed to estimate extinction risk, are often used to explore global patterns of diversity (Albuquerque & Beier, 2015; Strassburg *et al*., 2020; Allan *et al*., 2022). For plants, only ∼20% of the known species have been formally assessed at the global scale and sampling is non-random, with a particular focus on specific families or growth forms such as trees (Cannon *et al*., 2023). These limitations render IUCN range maps unsuitable for mapping global diversity patterns in plants and have contributed to the exclusion of plants in studies aiming to determine global conservation priorities, such as the minimum area of land required for the conservation of biodiversity or to determine priority areas for ecosystem restoration (Strassburg *et al*., 2020; Allan *et al*., 2022).

Open biodiversity portals such as the Global Biodiversity Information Facility (GBIF) hold hundreds of millions of digitally available plant occurrence records (Heberling *et al*., 2021), which can potentially be used to produce global maps of plant diversity (e.g., Daru, 2024). Such plant species occurrence data are, however, largely incomplete and biased taxonomically, temporally, and geographically (Meyer *et al*., 2016; Troudet *et al*., 2017). This means that global studies of plant diversity are limited to strongly biased occurrence data of a subset of known species in which the missing species are non-randomly distributed in their taxonomic and functional affinity or geographic distribution (Meyer *et al*., 2016; Cornwell *et al*., 2019; Boyd *et al*., 2023; Ondo *et al*., 2024). In fact, Ondo *et al*. (2024) revealed several “darkspots” of plant diversity where incomplete and biased digital knowledge of the spatial distribution of plants likely lead to misinterpretations of diversity patterns. Hence, it seems that existing fine-scale plant diversity maps are based on uneven and strongly biased digitally available plant occurrence data where, globally, certain groups and areas remain over- or under-represented (Meyer *et al*., 2016; Troudet *et al*., 2017; Cornwell *et al*., 2019; García-Roselló *et al*., 2023).

To promote the inclusion of plants in data-driven conservation efforts, it is thus necessary to find alternatives to better map diversity patterns from incomplete data (Brummitt *et al*., 2021). Studies aiming to produce maps of global plant diversity have used several approaches to overcome the existing limitations. The most common way to extrapolate beyond existing distribution data and create global diversity estimates, particularly for species richness, has been through species distribution models (SDMs) and machine learning methods (Brummitt *et al*., 2021; Jung *et al*., 2021; Sabatini *et al*., 2022; Qian *et al*., 2023). For example, Jung *et al*. (2021) included available occurrence data for ∼41% of the world’s vascular plants to model synergies between areas of conservation value for biodiversity, carbon and freshwater. More recently, Daru (2024) used SDMs to predict global patterns of vascular plant diversity for 201,681 vascular plant species (∼57% of all known vascular plant species). However, while SDM approaches can be used to predict diversity patterns based on incomplete distribution data, these methods are highly dependent on input data quality and quantity, and are likely to amplify taxonomic, geographic and environmental biases present in the input distribution data (Beck *et al*., 2014). Similarly, machine learning methods such as boosted regression trees have been used to model global plant diversity patterns based on vegetation plot data (Sabatini et al., 2022). While primarily used as an ecological and monitoring resource, large numbers of plant occurrence records across ecosystems are held in vegetation plots (e.g., sPlot (Sabatini *et al*., 2021)) or ForestPlots (ForestPlots.net *et al*., 2021)). The data from these plots are, however, heavily biased in terms of species composition because most vegetation plots in the tropics only include data from large trees, as well as geographically, because much more data is available for Europe than for the tropics.

Global taxonomic checklists such as the World Checklist of Vascular Plants (WCVP, https://powo.science.kew.org/; Govaerts *et al*. (2021)) provide a reliable reference of coarse plant diversity patterns at a global scale and have been used to map vascular plant distribution at a botanical country scale in several studies (Maitner *et al*., 2023; Tietje *et al*., 2023; Bachman *et al*., 2024; Ondo *et al*., 2024; Tordoni *et al*., 2024). Botanical countries largely correspond to global administrative areas, but several regions have been aggregated or further subdivided based on their phytogeographic differences (e.g., the Canary Islands are separated from mainland Spain due to their floristic distinctiveness, see Brummitt *et al*. 2001). The use of taxonomic checklists has the advantage that they are verified by experts and that all known species are listed even if there is insufficient knowledge of their range at finer scales in any given country. Recently, Cai *et al*. (2023) have used a near-complete global taxonomic checklist of plant distributions to predict diversity patterns at finer scales through machine learning approaches. While these maps likely suffer less from the biases concerning occurrence data, (in terms of their taxonomic and geographic coverage), predictions at finer scales require smoothing and extrapolation beyond the resolution of the existing data, which comes with its own caveats.

An alternative but little-explored approach that has the potential to combine the strengths of taxonomic checklists with fine-scale distribution data would be representative sampling, where sampling is guided by verifiable patterns of diversity (e.g., a global taxonomic checklist at botanical country scale for plants) to create a robust dataset for a smaller subset of species that is representative of our current knowledge of plant diversity (Boyd *et al*., 2023, 2024). Approaches based on a smaller samples of species are already used in conservation and ecology to generalise patterns across groups, such as in environmental monitoring (Boyd *et al*., 2023) or to estimate extinction risks (Baillie *et al*., 2008; Brummitt *et al*., 2015). While representative sampling has obvious disadvantages such as the loss of information from excluded species (Boyd *et al*., 2023), representative sampling can help to (1) reduce biases, both in terms of taxonomic and geographic affinities of the species, (2) allow for better quality control as data gaps can be more easily prioritised and consolidated, and (3) enable the incorporation of prior knowledge about diversity for consistent mapping of patterns (Boyd *et al*., 2023).

Sample-based approaches to mapping global diversity patterns remain largely untested for vascular plants. A previous study focused on identifying families as model groups representative of species richness patterns for vascular plants globally (Nic Lughadha *et al*., 2005), with Fabaceae emerging as the candidate model family. While choosing one (or several) plant groups as a surrogate for all plants has advantages (e.g., improved data quality and taxonomic stability), this kind of approach does not capture the full spectrum of ecological strategies and evolutionary history of other plant species, which would be better done through a sampling approach that can be measured through a range of diversity metrics. Additionally, sampling approaches potentially require a much smaller number of species to achieve patterns representative of all plants (Baillie *et al*., 2008; Brummitt *et al*., 2015). For example, fewer than 2% of all plant species were used to create the Sampled Red List Index (Brummitt et al. 2015) which aims to quantify threats for all plant species based on a random sample of species. A previous attempt of using distribution data from a random sample of species to model global species richness by Brummitt *et al*. (2021) also shows starkly different species richness patterns compared to maps produced from all available data (Jung *et al*., 2021; Daru, 2024), suggesting that either one or the other approach to mapping plant diversity might indeed distort global species richness patterns.

A first step towards testing the robustness of a representative sampling approach and ultimately creating a dataset that best represents our current knowledge of plant diversity patterns would be to determine the number of species needed to reliably re-create diversity patterns at the finest scale where a comprehensive reference dataset is available. The results of such a study could then be used to translate broad patterns to finer spatial scales. This is particularly the case for dimensions of diversity beyond species richness such as phylogenetic diversity and measures that incorporate species community properties where thresholds have not formally been tested before. Here we establish and test such a representative sampling method for vascular plants by identifying the minimum number of species needed to recreate different aspects of global vascular plant diversity based on random sampling, using the WCVP as our reference dataset for known vascular plant diversity patterns. Specifically, we aim to (1) test the feasibility of representative random sampling to accurately map global diversity patterns and how this performs when compared against other taxonomic groupings (families, genera), (2) provide first estimates of sample sizes needed to achieve robust predictions, and (3) provide a baseline for the creation of a dataset based on a representative subset of species that enables robust predictions of different measures of plant diversity at a global scale. Determining this number of species is a crucial step towards the mapping of fine-scale patterns of plant diversity that is needed for accurate conservation prioritisation, and it has important implications for conclusions drawn from previous studies of plant diversity patterns.

## Materials and Methods

We used the WCVP (version 12) to determine reference diversity patterns within 368 botanical countries (Govaerts *et al*., 2021). The WCVP is an expert-curated checklist that includes data on the status of plant names; each accepted species name is associated with verified distribution data at the scale of a botanical country from the World Geographical Scheme for Recording Plant Distributions (WGSRPD) (Brummitt *et al*., 2001). We kept only accepted names at species rank, excluded hybrid species and species considered extinct, and only retained distribution data for countries where species were considered to be native, and where records were not labelled as doubtful. The final dataset amounted to 349,113 accepted species in 13,899 genera and 454 families across 368 botanical countries amounting to 1.3 million distribution records used for our analysis.

### Phylogenetic and growth form data

To estimate a phylogeny of all vascular plants, we used a publicly available phylogenetic backbone (Jin & Qian, 2022a) to construct a phylogeny of all plant species included in our dataset via the U.phylomaker package in R using Scenario 3 (see Jin & Qian (2022b) for more information). We inserted species missing from the phylogeny at genus or family level using the finest taxonomic level available in the phylogeny, including 21,956 species inserted based on family, 258,126 species inserted based on genus, and 69,031 species that were found in the megaphylogeny.

We used growth form as a measure of functional diversity, as it captures many aspects of plant life in a single character and is available for most plants. We determined the growth form of each species using publicly available standardised data from the GIFT database (Weigelt *et al*., 2020; Taylor *et al*., 2023). We chose this over the data available from WCVP as plant growth forms were already standardised, data were available for a higher proportion of species, and the databases use the same taxonomic backbone. For species with multiple recorded growth forms, all species that contained epiphyte, climber or herb in combination with another growth form were designated as epiphyte or climber or herbs, respectively, in our dataset. For other species with multiple growth forms (herb, subshrub, shrub, tree) a hierarchical system to assigning growth forms based on plant height (subshrub (0.5m) < shrub (0.50–10Lm) < tree (>L10Lm)) was used. This provided growth form data for 271,509 species (∼78 % of all species used). For ∼22% species where no or only ambiguous growth form data was obtained, we used the “nnet” package in R (Venables & Ripley, 2002) to run a multinominal logistic regression to predict their growth form using plant family, major growth form of other species in the genus and geographic information (TDWG level 2 - regional scale) as predictor variables. For genera lacking growth form altogether, we ran the model without major growth form as a variable. Based on the original data we achieved accuracy of >0.8 for the model including all predictor variables, which was used to impute growth forms for 75,132 species, whereas the model including only family and TDWG level 2 information achieved an accuracy of 0.67 and was used for 2,472 species (Table SI1). Our results closely matched predictions from previous studies suggesting that herbs are the most common growth form while trees and shrubs make up similar proportions of all species, respectively (König *et al*., 2019; Kattge *et al*., 2020) and also matched the number of epiphytic species from Zotz *et al*. (2021).

### Analysis

The first step in our analysis consisted of creating reference diversity maps based on four diversity metrics of plant diversity commonly used or already employed in similar studies: (1) species richness, (2) phylogenetic diversity, (3) effective family and (4) growth form diversity. Species richness (the number of species for each botanical country) is the most frequently applied diversity metric for conservation purposes and the one most readily available for large parts of the world. We calculated phylogenetic diversity as the sum of branch lengths (Faith’s PD) covered by the species of each botanical country using the “phyloregion” package (Daru *et al*., 2020). We chose this measure to represent the diversity of evolutionary histories in each botanical country. Both species and phylogenetic diversity are incidence-based measures that do not incorporate measures of community similarity. To address this, we also used the number of species per plant family and growth form in each botanical country to capture aspects of floristic and ecological diversity in their communities, respectively. We calculated these diversity indices using Hill numbers, also referred to as the effective number of species (Hill, 1973; Jost, 2006, 2007; Chao *et al*., 2014), with *q* = 1. Hill numbers are a measure of diversity that integrate both group frequencies and richness, with the sensitivity of the metric to either rare or dominant groups determined by the value of the parameter *q* (Chao *et al*., 2014). The Hill number framework provides a standardised framework of calculating diversity indices and has become an increasingly common way of presenting diversity estimates (Chao *et al*., 2014; Chiu *et al*., 2014; Luypaert *et al*., 2022; Byrnes *et al*., 2023; Dick, 2023); for growth forms, a similar approach has also recently been taken by Taylor *et al*. (2023) who measured the diversity of growth forms at a global scale as the effective number of growth forms. At *q* = 1 this measure equates to the exponential of the Shannon Index, also referred to as Shannon diversity, and emphasises neither common nor rare species (Chao *et al*., 2014) which is why we chose this value for our analysis as it treats groups equally. The resulting value can thus be interpreted as the “effective number of groups” (i.e., the number of equally frequent groups that would yield the observed Shannon diversity). We therefore refer to the metrics applied here as “effective family diversity” and “effective growth form diversity”.

To determine the number of species needed to re-create the reference diversity patterns, we used random sampling from the global species pool of vascular plants to generate samples for sample sizes from 1 - 349,113 species iteratively increasing the number of species included in the sample by one for each diversity metric. At each sample size we created 100 independent random samples taken without replacement. We used the R package “GWmodel” (Lu *et al*., 2014; Gollini *et al*., 2015) to calculate local spatially weighted Spearman correlation coefficients to determine the correlation between the spatial pattern of diversity in each sample and the reference diversity pattern based on all species. We chose this approach because we detected significant spatial autocorrelation in the distribution data (Moran’s I: 0.55), which can lead to inflated correlation coefficients (Osorio *et al*., 2020). Spatially weighted Spearman correlation coefficients use a bi-square kernel function to calculate local correlation coefficients for each botanical country based on a set of nearest neighbours, where weight of neighbouring botanical countries decays with distance (distances were based on the centroid of each botanical country). We chose the number of nearest neighbours based on an automated leave-one-out cross-validation approach implemented in the “GWmodel” package. We calculated the median of the local correlation coefficients, hereafter referred to as the global correlation coefficient.

For each sample size we calculated the mean and the confidence interval of the global correlation coefficient, which enabled us to determine the minimum number of species needed to reach a correlation of ≥ 0.95. We determined ρ ≥ 0.95 as our threshold for representativeness to mitigate the sensitivity of the calculations to outliers (instead of a higher threshold) and to provide an analogy to align with conventionally employed error-tolerance boundaries. We then applied Kruskal-Wallis and *post-hoc* Dunn’s tests to identify significant differences between the number of species included to reach a global correlation coefficient of 0.95 for different diversity metrics. To further compare diversity values at different sample sizes, we extracted values of each metric from 100 random samples of the lowest (1,048 *spp*.) and highest (67,772 *spp*.)mean number of species needed to meet the threshold among the different diversity metrics. We then produced maps using min-max normalisation to achieve a common scale and compare diversity patterns visually. In min-max scaling the raw diversity values are rescaled using the range of the variable (x_norm_ = x − x_min_ / x_max_ − x_min_), thereby bringing different variables to the same scale with values between 0 and 1, while preserving the original distribution of the values. We formally analysed differences between the datasets by comparing the average ranks of botanical countries using the deviation of the mean rank in the sample from the original ranks (x_dev_ = x_mean_ – x_org_) as a measure of representativeness. We calculated the coefficient of variation (cv = (standard deviation / mean) * 100) to estimate the variation in diversity estimates across the samples.

To compare the representativeness of plant families and genera against the results of random sampling, we calculated global correlation coefficients for the species richness patterns of each plant family and genus, comparing them against (1) the reference species richness patterns with all species included and (2) the average global correlation coefficient of 100 random samples at the sample size corresponding to the number of species in the plant group examined.

## Results

### Thresholds of representativeness

The speed of increase of the correlation between the diversity patterns of a random sample of species compared to the global reference patterns of diversity for all species differed between diversity metrics (Fig. 1A). Incidence-based metrics showed similar patterns in reaching the threshold of a global correlation coefficient of 0.95 to between 1,048 species (Standard error (SE)=9, species richness) and 1,579 species (SE=15, phylogenetic diversity) on average (Fig. 1B, Table S1). In contrast, metrics that incorporated species frequencies for different groups reached the threshold much later on average, at 35,127 species (SE=403, effective growth form diversity) and 67,772 species (SE=235, effective family diversity; Fig. 1B, Table S1).

**Fig. 1.**
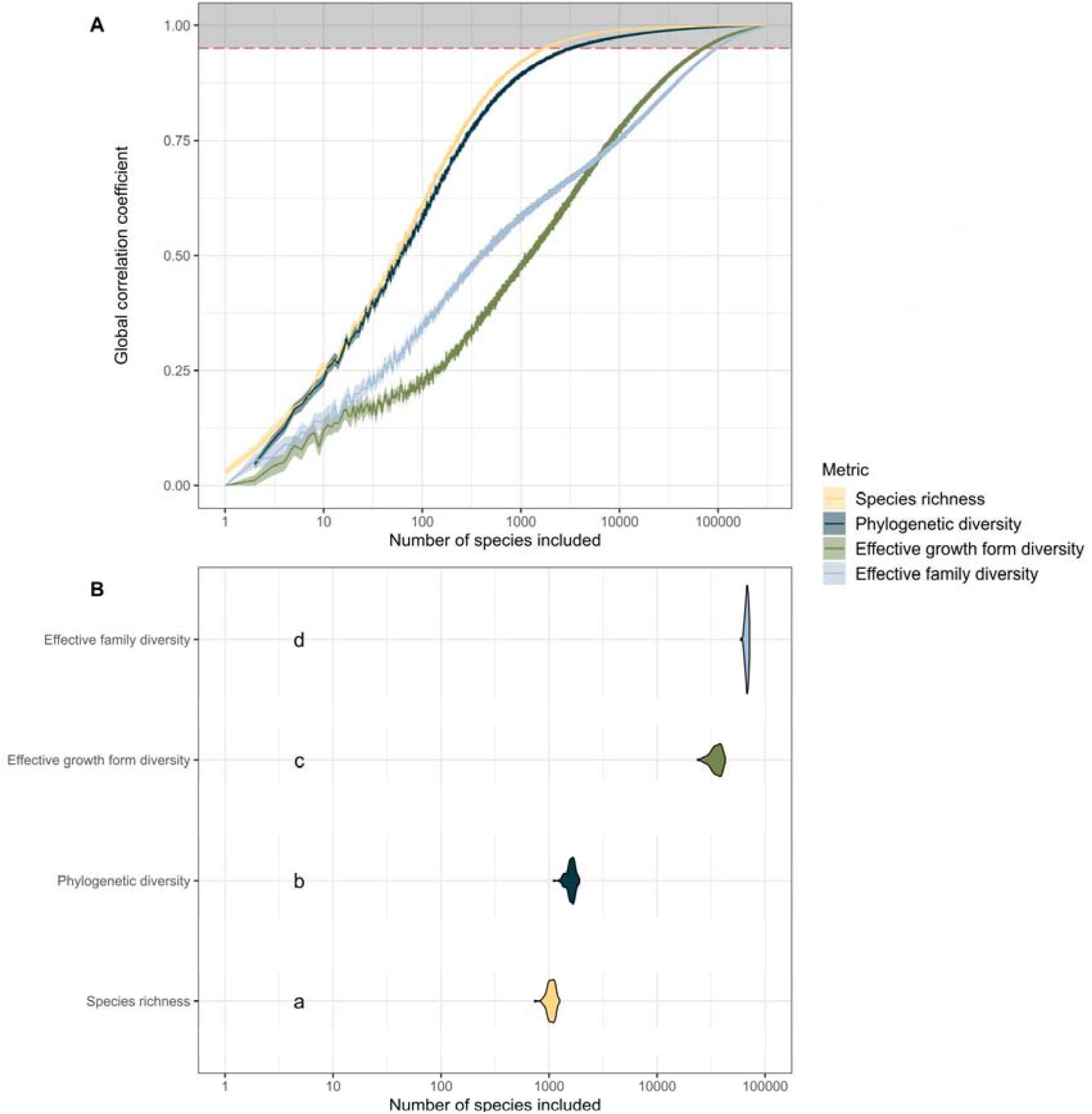
A) Accumulation curves of the average global correlation coefficients of four diversity metrics across 100 random samples when comparing their diversity patterns to the reference diversity patterns based on all vascular plant species. Shades around the lines represent 95% confidence intervals of the mean. The grey zone marks where a group is considered representative i.e. the correlation is ≥ 0.95. The x axis is on a logarithmic scale (base = 10). B) The number of species needed in order to achieve a global correlation coefficient of 0.95 in each of the 100 samples. Kruskal-Wallis and post-hoc Dunn’s tests were used to identify significant differences between the number of species needed to meet the threshold for each diversity metric. Letters indicate significant differences between the groups.

For the species richness metric, we also tested the strength of the correlation between the distribution of species richness for each vascular plant family and genus and global reference species richness patterns. No individual family or genus met the threshold of 0.95. Some families and genera outperformed random samples at lower sample sizes. For families, the highest sample size where a family performed better was 161 species (Hypoxidaceae, global ρ = 0.68, Fig. 2A), whereas for genera the number of species was 135 (*Setaria,* global ρ = 0.70, Fig. 2B). The top five families that followed the global species richness pattern the best were the Fabaceae (global ρ = 0.94, n = 22,235), the Poaceae (global ρ = 0.93, n = 11,801), the Lamiaceae (global ρ = 0.92, n = 7,696), the Asteracae (global ρ = 0.91, n = 34,138), and the Euphorbiaceae (global ρ = 0.90, n = 6,495; Fig. 2A). The top five genera that followed the global species richness the best were *Selaginella* (global ρ = 0.81, n = 727), *Cyperus* (global ρ = 0.81, n = 950), *Polygala* (global ρ = 0.80, n = 668), *Euphorbia* (global ρ = 0.80, n = 2,038), and *Clematis* (global ρ = 0.80, n = 385, Fig. 2B).

**Fig. 2.**
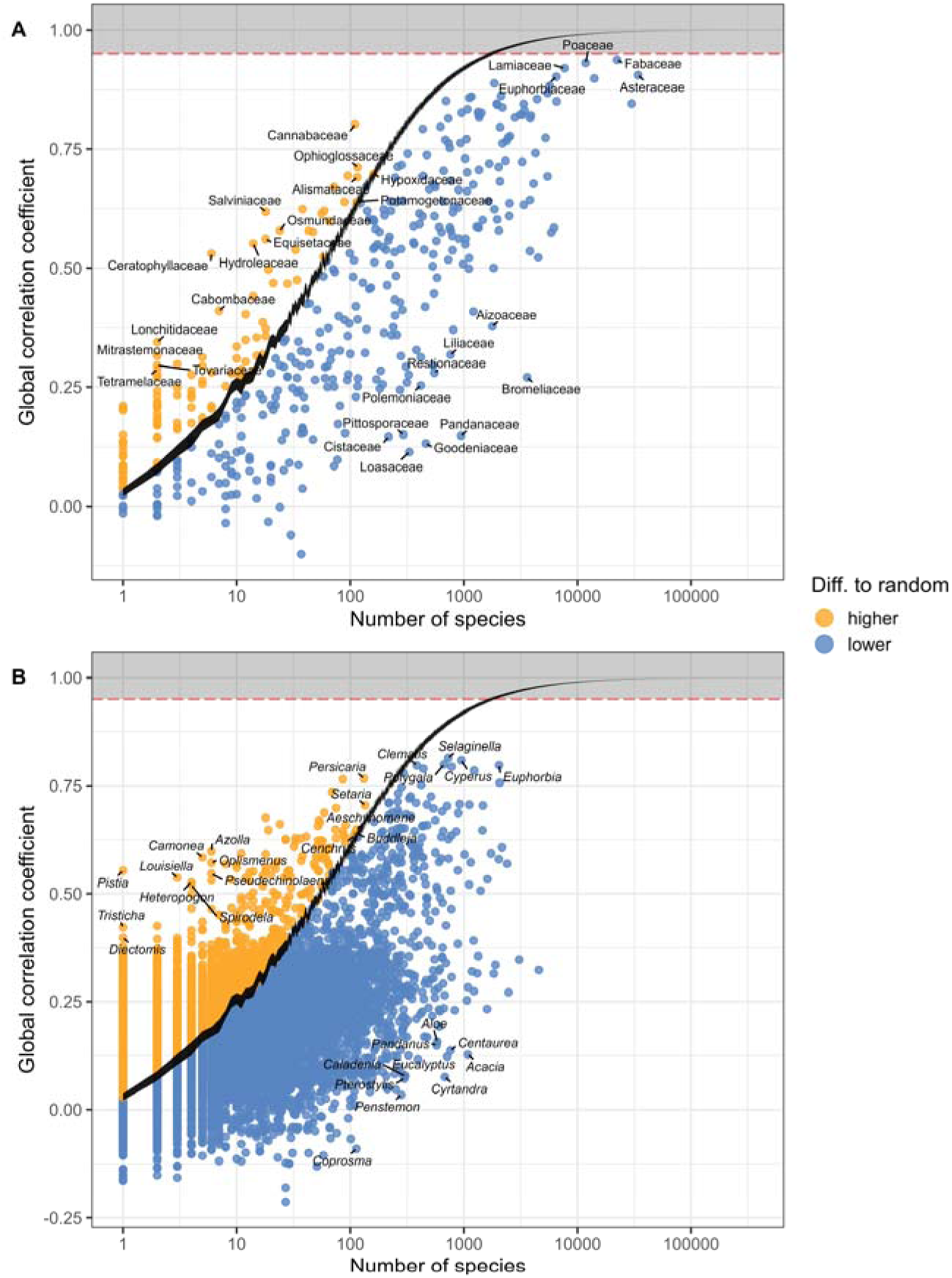
Global correlation coefficients of species grouped by A) plant families and B) plant genera when compared to the reference patterns of species richness based on all vascular plant species. The black line marks the confidence interval of the global correlation coefficient based on 100 random samples with iteratively increasing sample sizes. Orange dots have higher correlations than the random sample, while blue dots have lower global correlation coefficients. The grey zone marks where a group is considered representative i.e. the correlation is ≥ 0.95. The x axis is in a logarithmic scale (base = 10)

### Diversity patterns

Our results are generally in line with previous vascular plant diversity maps (Tietje *et al*., 2023) in showing both the highest species richness and phylogenetic diversity in Colombia (*Fig. 3*A, D). Effective growth form diversity (i.e. the number of equally frequent growth form groups to observe this level of diversity) was highest in Brazil Southeast (*Fig. 3*G). For effective family diversity, China Southeast had the highest diversity values (Fig. 3J). Across all diversity metrics, South Sandwich Islands showed the lowest values.

**Fig. 3.**
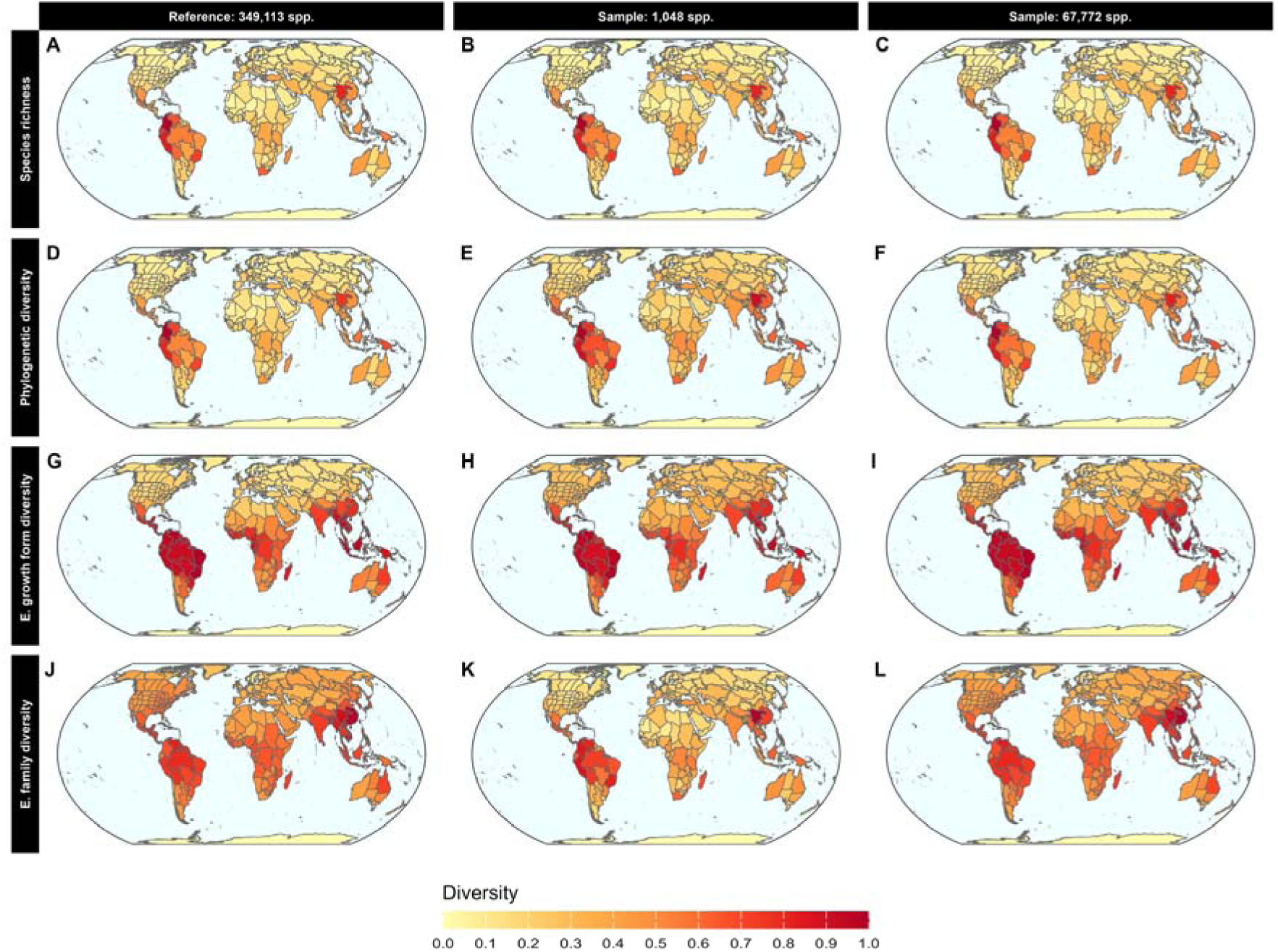
Diversity maps based on the full reference data (all vascular plants; left column) and the average diversity at the smallest average (middle column) and largest average (right column) number of species needed to achieve a global correlation coefficient of ≥ 0.95 across 100 random samples. Each row corresponds to a different diversity metric (species richness, phylogenetic diversity, effective family and effective growth form diversity). Diversity values are presented using a min-max normalisation to achieve a common and comparable scale. The map is displayed using the equal earth projection.

Diversity maps recovered from the lowest and highest sample size needed to meet diversity metric thresholds closely resembled the original global patterns (Fig. 3). Particularly for species and phylogenetic diversity, average rank differences were low across all continents in both samples (Fig. 4). In the smaller sample of 1,048 both the effective family and effective growth form diversity of botanical countries in Europe or Asia-Temperate was on average over-estimated, while diversity metrics of botanical countries in Asia-Tropical, the Pacific and Southern America were on average under-estimated (Fig. 3, Fig. 4). For botanical countries in Northern America, effective growth form diversity was over-estimated while effective family diversity was under-estimated (Fig. 3, Fig. 4H). In the largest sample of 67,772 species, only botanical countries in the Pacific and Antarctica were under-estimated in terms of their effective growth form diversity (Fig. 3, Fig. 4C, I). Botanical countries in the Pacific were also slightly under-estimated in terms of their effective family diversity (Fig. 3, Fig. 4J). The effective family diversity of botanical countries in Asia-Temperate and Europe was significantly over-estimated in the largest sample of 67,772 species (Fig. 3, Fig. 4D, G).

**Fig. 4.**
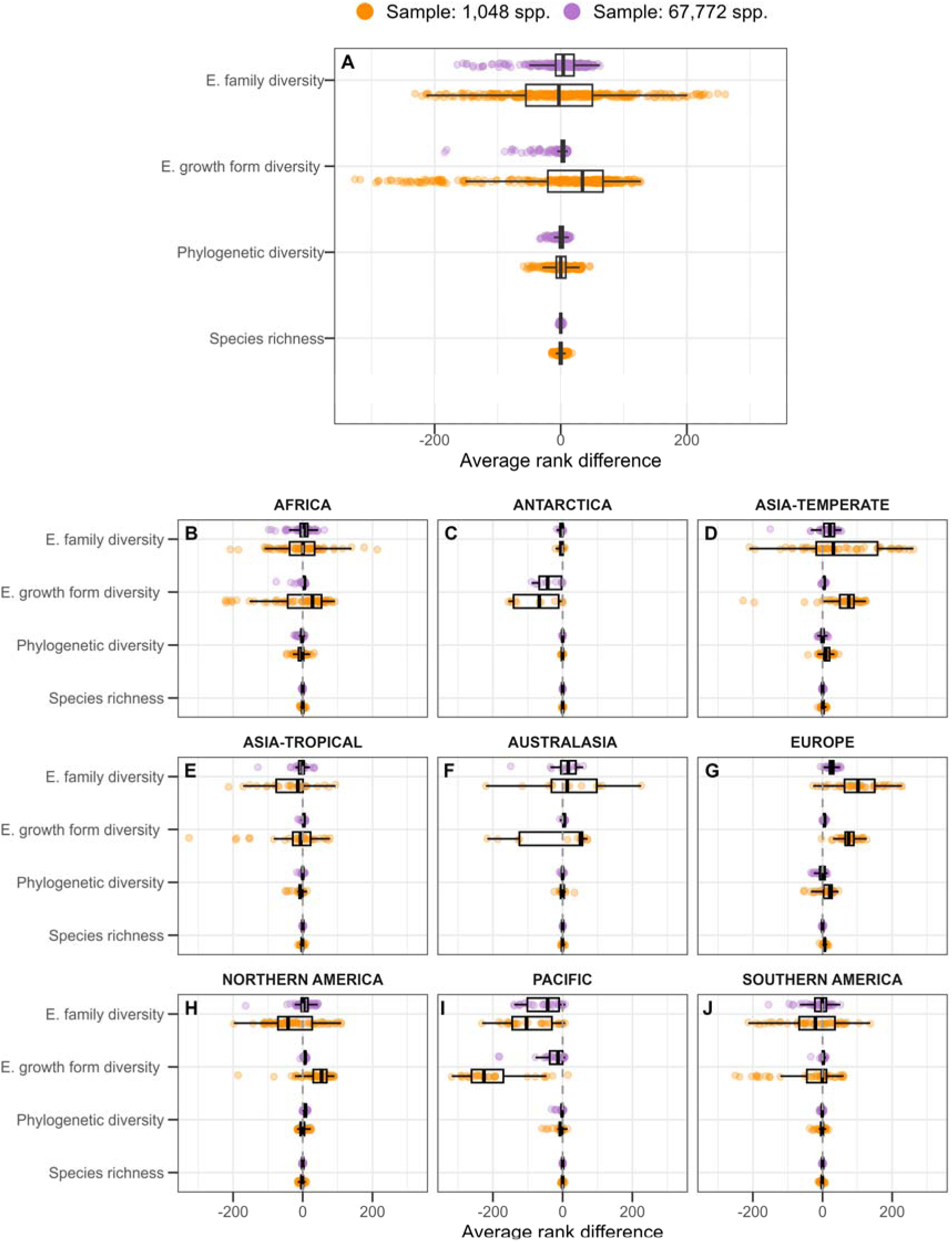
Rank difference of botanical countries between their average rank across 100 random samples and their original rank in the reference data when including all vascular plant species for the smallest average (orange) and largest average (purple) sample size needed to achieve a global correlation coefficient of ≥ 0.95. The plots show the data for A) all botanical countries included in the analysis and B-J) botanical countries split by continents.

Variation within the 100 random samples, as measured by coefficients of variation (ratio of standard deviation and mean, CV), was relatively low across all four diversity metrics when sampling 67,772 species (average CV of 6.9-9.2%, Fig. 5, Table SI3) but considerably higher when sampling only 1,048 species (average of CV 58.0-67.2%, Fig. 4, Table SI3). The CV was also strongly correlated to the average diversity rank of each botanical country, with botanical countries with less diversity having the highest CV values (Figure SI1). Average rank differences were relatively high for some metrics, with the smaller samples having 1,048 species (3.4-75.2, Table SI3), and were much reduced with the largest sample of 67,772 species (0.5-23.8; Fig. 4, Table SI3). The maximum rank differences with the smallest sample of 1,048 species were 18 for species richness, 55 for phylogenetic diversity, 327 for effective growth form diversity and 261 for effective family diversity (Fig. 4, Table SI3). The maximum rank differences with the largest sample of 67,772 species were 5 for species richness, 33 for phylogenetic diversity, 184 for effective growth form diversity and 164 for effective family diversity (Fig. 4, Table SI3).

**Fig. 5.**
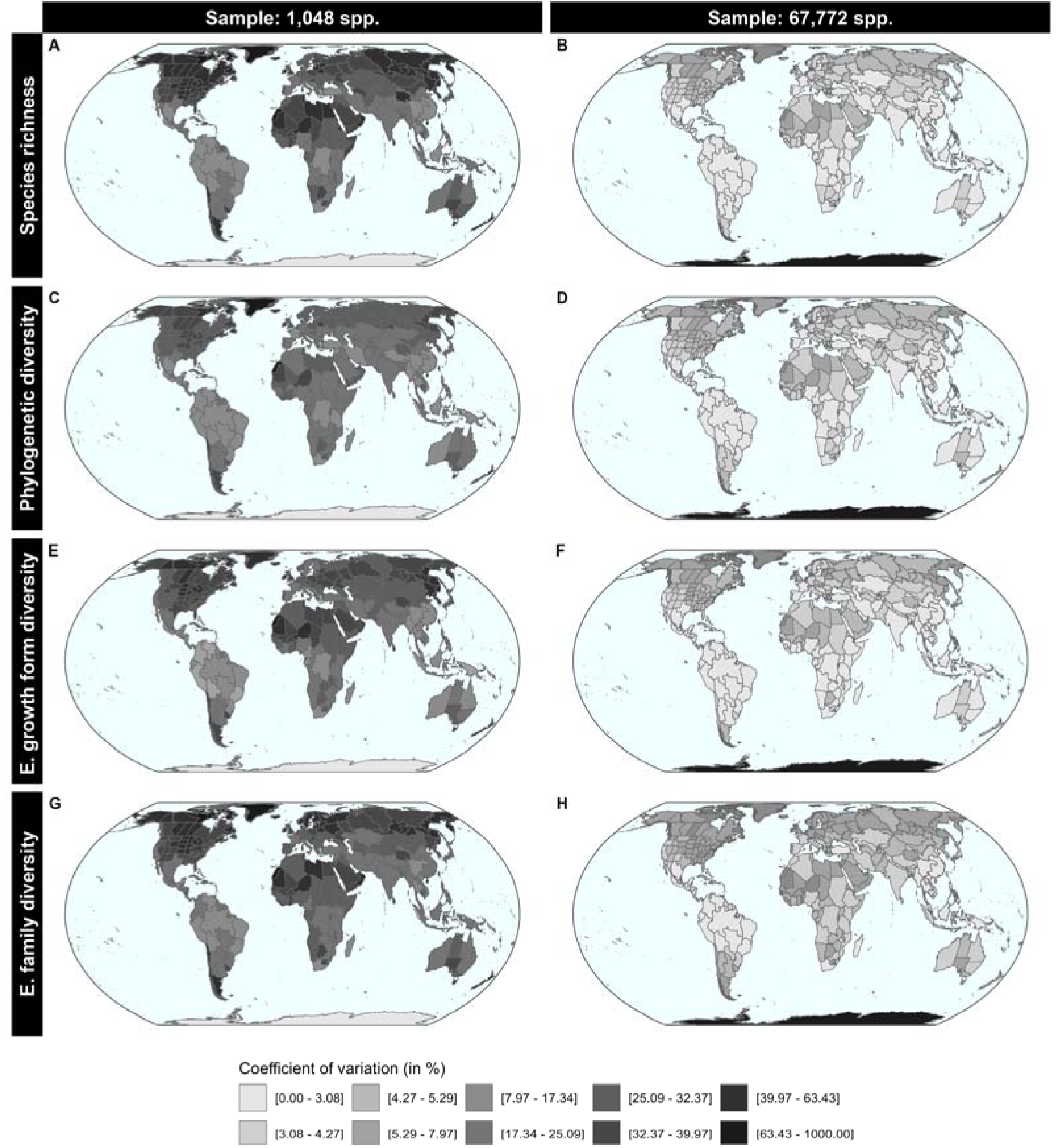
Variability maps of the different diversity metrics based on the average difference of each botanical country in 100 random samples from the reference diversity values at the smallest average (left column) and largest average (right column) sample size needed to achieve a global correlation coefficient of ≥ 0.95. The CV is the ratio of the standard deviation to the mean and is displayed as a percentage. The map is displayed using the equal earth projection, with colours based on data deciles.

## Discussion

Plants often remain poorly represented in global studies, including those for conservation prioritisation, in part because reliable maps that accurately display patterns of diversity for such a speciose group of organisms are still lacking. Here, we tested whether a sampling approach can yield representative and stable maps of the global patterns of multiple aspects of plant diversity at the finest scale where comprehensive data is available, using taxonomic checklists and random sampling. We show that, at the scale of botanical countries, accurate and representative maps of global vascular plant diversity patterns can be created from small, representative subsets of randomly selected species. For this study we measured four relevant biodiversity metrics with which we aimed to cover the multi-dimensionality of plant diversity. Only 0.4% of the total species (1,579, Fig. 1, Table SI2) were needed to map species and phylogenetic diversity, two of the most commonly used diversity metrics, accurately. For metrics that incorporated both group-richness and evenness, the number of species needed was a lot higher (67,772, 19.4%, Fig.1, Table SI2) but still fewer than 20% of total species were needed to create highly representative maps of effective growth form and family diversity at the scale of botanical countries. Using representative samples thus enables accurate inferences about global patterns using only a fraction of all species.

Species richness is the most commonly applied and universally used measure to determine areas of high conservation value and has high relevance for conservation planning (Fleishman *et al*., 2006). Species richness does not, however, account for variation in shared evolutionary history, taxonomic affinities or functional traits between species (Faith, 1992; Cadotte *et al*., 2011; Chao *et al*., 2014; Tordoni *et al*., 2024). We therefore included Faith’s phylogenetic diversity as an alternative measure to consider the feature diversity of the species included in the sample (Faith, 1992). To further extend the range of metrics we also examined how well this sampling approach captured the frequencies of families and growth forms within each botanical country using Hill numbers (Hill, 1973). Hill numbers have become one of the most common ways to unify different diversity metrics (Hill, 1973; Jost, 2006; Chao *et al*., 2014). We chose to include these metrics as they reflect the number of functional groups in the sample and how diversity is shared among them (evenness), which is an important aspect of diversity (i.e. one might interpret a system as more diverse for conservation purposes if all groups are equally frequent, rather than dominated by one group). These results suggest that higher numbers of species are needed to reflect community evenness across groupings: species and phylogenetic diversity reached the threshold at very low sample sizes (less than 1% of the species), whereas effective growth form and effective family diversity needed higher sample sizes to reach the same thresholds (∼10% and ∼19% respectively, Fig.1, Table SI1).

Our results also highlight that no single plant family or genus can capture the distribution of global plant diversity patterns as effectively as a random sample. While there are several benefits of using model plant groups to study general plant diversity patterns (Nic Lughadha *et al*., 2005), such as a consistent knowledge of the groups’ taxonomy and taxonomically verified distribution data and the availability of comprehensive phylogenies, diversity patterns of a single plant genus or family are less well-suited to capturing the whole spectrum of ecological, phylogenetic and functional diversity of all plant species compared to a random representative sample across all vascular plants. Combining several groups can overcome these limitations; however, representative sampling yields more robust results while requiring considerably fewer species (Fig 2). Focusing on a representative sample of species therefore harnesses the same benefits as focusing on specific plant groups, while providing deeper insights about the different aspects of plant diversity.

These results are not independent from choice of metrics, and we would expect different diversity metrics to require different but similar minimum sample sizes to reach our threshold of representativeness. Another common way to measure diversity, which was not included here, is through functional diversity metrics that aim to measure the diversity of species traits through distance-based or hypervolume-based approaches (Cadotte *et al*., 2011; Mammola *et al*., 2021). Currently there is no comprehensive trait dataset with enough data to enable tests across all vascular plant species (Maitner *et al*., 2023), so to enable a realistic estimate of the number of species needed to cover the functional dimension of plant diversity at the global scale, these asymmetries in the availability of trait data have to be addressed. This would be particularly interesting as a recent study has shown that in plants functional and phylogenetic diversity are likely decoupled (Hähn *et al*., 2025). We also focused on alpha diversity values and did not test the influence of sample size on beta-diversity metrics that take into account the variation of species between sites (Anderson *et al*., 2011). While alpha diversity is useful to capture absolute peaks in diversity, beta diversity metrics can have important implications for conservation about the regional organisation of diversity, such as identifying areas with particularly distinct species assemblages, and conserving regional diversity (Socolar *et al*., 2016). Including functional diversity and beta diversity metrics would thus be an important next step for future studies.

While our study underlines the robustness of previous approaches for conservation based on a similar methodology (e.g. the Red List Index and the Sampled Red List index, following Baillie et al., 2008; Brummitt et al., 2015), adapting this framework to support detailed spatial conservation planning will require further steps. Firstly, we sampled at random to generate representative subsets of our reference dataset (with the assumption that this was largely unbiased); however, this approach is unlikely to be feasible for other datasets needed to model species distributions. In practice, most datasets used to date for mapping or modelling the spatial distribution of plant diversity are unrepresentative (Meyer *et al*., 2016; Boyd *et al*., 2023), meaning large parts of the phylogenetic and taxonomic diversity of plants are excluded from analyses (Cornwell *et al*., 2019). Extracting a random subset of species from such datasets would therefore merely yield a biased sample. To achieve representative sampling, optimisation algorithms or machine learning methods could be used to enhance the representativeness of the sample from a random starting point by iteratively improving the properties of the species included in the sample when compared to the reference patterns (Tominaga, 1998; Kamp & Savenije, 2006; Akdemir *et al*., 2015; Browning *et al*., 2017; Zhang & Zhu, 2019; Hauptmann *et al*., 2023). Employing these methods would drastically decrease the number of species needed to reach a high level of representativeness, as has already been shown for the number of sites needed to design efficient environmental monitoring systems (O’Hare *et al*., 2020).

Secondly, the analysis presented here was conducted at the botanical country scale, which currently corresponds the finest spatial resolution where comprehensive plant distribution data are available (Brummitt *et al*., 2021). However, this spatial scale remains far too coarse to fully inform detailed conservation and restoration planning (Cohen & Jetz, 2025). Also, the sample size estimations presented here are influenced by the number and size of the units of the distribution data used in the analysis and as such are scale dependent (Jarzyna & Jetz, 2018). As global consensus checklists and range maps at even finer spatial scales become available, we would expect more species to be needed to reach the same level of representativeness, with the patterns increasing in complexity as sampling units are added. Further analyses are therefore needed to establish the number of species required to achieve robust predictions of plant diversity patterns with a representative subsample at fine spatial scales.

Simulation-based methods offer a promising opportunity to determine these finer-scale thresholds. Studies have shown that macroecological patterns (range size, abundance, taxonomy) are often log-normally distributed (Gaston, 1996; Bell, 2001; Frodin, 2004). Globally, most plant species are rare, while relatively few are common (Enquist *et al*., 2019); equally, only a few areas harbour most of the species (Mittermeier *et al*., 2011), such as mountains that make up ∼25% of the land area but house 85% of all mammal, amphibian and bird species (Rahbek *et al*., 2019). The lognormal distribution of macroecological patterns could therefore be used and supplemented with additional environmental data to create virtual species based on known properties of ecological data and responses of plants to environmental conditions (reviewed by Miller, 2014). Virtual species could then be used to map patterns of diversity at different sample sizes without the need for fine-scale distribution data, and to estimate the number of species at which diversity metrics for a given spatial resolution no longer change significantly. Similarly, sample-size thresholds could be determined by repeatedly modelling sets of species, based on existing fine-scale distribution data, with varying sample sizes that are representative at the botanical country scale.

Ultimately, to create a robust, fine-scale dataset for a representative subset of species, taxonomically verified and georeferenced records of plants are needed (Meyer et al., 2016). Studies of data collection patterns in animals have already shown data to be biased towards areas with higher population densities and near roads (Hughes *et al*., 2021). Similar biases are evident in digitally available plant occurrence datasets (Daru *et al*., 2018; Daru & Rodriguez, 2023). While big data approaches often do not allow for detailed data exploration and rely on standardised data-cleaning processes, using a smaller sample has the advantage that data gaps can be closed more easily with the help of experts, and additional data sources such as information from floras and non-digitised herbarium collections can be more readily integrated. Delves et al. (2023) have already shown that incorporating collections from small herbaria that are currently not incorporated within global databases can greatly improve the accuracy of range and threat estimates of species, particularly for rare species. Rare species are often overlooked in large-scale analyses due to data gaps (Jeliazkov *et al*., 2022) but in a small, representative sample could be more readily included through targeted data collection and curation efforts, thus enhancing their proportional and spatial representation. Focusing on a representative subset of species could therefore help the botanical community in terms of where to focus practical conservation efforts.

## Conclusion

As we head into a decisive decade for biodiversity, detailed patterns of plant diversity and their drivers are still poorly understood (Tordoni et al., 2024). Equally, our ability to produce representative, data-driven, global maps of plant diversity patterns at high resolution to be used for conservation planning on the ground remains limited. Closing these knowledge gaps for all species would take decades (Ondo et al., 2024). It is therefore necessary to focus on finding alternatives to estimate biodiversity patterns better from current, incomplete data. Here, we present an approach that is less data-intensive and can circumvent data gaps, while still enabling robust inferences about biodiversity patterns at a global scale. We also show that a representative sampling approach outperforms other approaches and allows us to (1) close remaining knowledge gaps faster, (2) facilitate both data curation and collection, and (3) address existing biases more directly than in large datasets. We thus hope that the approach presented here can serve as a basis for future studies aiming to create representative datasets in order to ensure that plants are considered appropriately in systematic conservation planning.

## Acknowledgements

LB’s work is supported by a NERC Doctoral Training Partnership grant (NE/S007407/1). We thank everyone involved in contributing to the ongoing efforts to document biodiversity globally.

## Author contributions

LB, PWM, NAB, SP, and TS planned and designed the research. LB conducted the data analysis and wrote the manuscript. PWM, NAB, SP, and TS helped to revise the manuscript.

## Data accessibility

The main raw data supporting this manuscript is freely available online at https://powo.science.kew.org/. Data supporting the main analysis of the manuscript are stored on OSF under https://doi.org/10.17605/OSF.IO/N8WUB. The R code to reproduce the main analysis presented in the manuscript is available from https://github.com/baldasztil/Sampling_wcvp.

