## Supporting Information for "Small representative samples can capture global vascular plant diversity patterns"

The Tables and figures in this document support the findings of:

### Tables

Table SI1 Model performance statistics for our multinominal model to predict growth forms of species with missing growth form information shown for two different model specifications. Model 1 takes into account the most common growth form in each species family, the region of occurrence and in the genus. Model 2 was used when the genus did not have any growth form information. Model accuracy was assessed based on their predictions for species with known growth forms. ppv = positive predictive value, npv = negative predictive value, mcc = Matthews correlation coefficient, j_index = Youden's J statistic.

| **Metric** | **Estimate** | **Model** |
| --- | --- | --- |
| **accuracy** | 0.818 | **model_family + major_region + major_growth** |
| **kappa** | 0.760 |  |
| **sensitivity** | 0.789 |  |
| **specificity** | 0.960 |  |
| **ppv** | 0.804 |  |
| **npv** | 0.962 |  |
| **mcc** | 0.762 |  |
| **j_index** | 0.749 |  |
| **balanced_accuracy** | 0.875 |  |
| **detection_prevalence** | 0.167 |  |
| **precision** | 0.804 |  |
| **recall** | 0.789 |  |
| **f_measure** | 0.791 |  |
| **accuracy** | 0.667 |  |
| **kappa** | 0.560 | **model_family + major_region** |
| **sensitivity** | 0.610 |  |
| **specificity** | 0.927 |  |
| **ppv** | 0.643 |  |
| **npv** | 0.929 |  |
| **mcc** | 0.561 |  |
| **j_index** | 0.537 |  |
| **balanced accuracy** | 0.768 |  |
| **detection_prevalence** | 0.167 |  |
| **precision** | 0.643 |  |
| **recall** | 0.610 |  |
| **f_measure** | 0.619 |  |

Table SI2 Summary statistics for the number of species needed to reach a geographically weighted global correlation coefficient of 0.95 when compared against the reference diversity patterns of vascular plants. Calculations are based on 100 random samples and thresholds are displayed for each diversity metric. Values are rounded up to integers. Min = minimum, Max = maximum, MAD = mean absolute deviation, SE = standard error, CI = Confidence interval

| Diversity metric | Mean | Min | Max | Median | MAD | SE | 95% CI |
| --- | --- | --- | --- | --- | --- | --- | --- |
| Species richness | 1,048 | 735 | 1,253 | 1,054 | 85 | 9 | [1,031, 1,065] |
| Phylogenetic diversity | 1,597 | 1,099 | 1,915 | 1,610 | 126 | 15 | [1,568, 1,625] |
| Effective growth form diversity | 35,127 | 23,607 | 42,815 | 35,524 | 4,580 | 403 | [34,337, 35,916] |
| Effective family diversity | 67,772 | 59,555 | 71,795 | 67,874 | 2,480 | 235 | [67,312, 68,232] |

Table SI3 Summary statistics for 100 random samples extracted at the lowest and highest number of species needed to reach a geographically weighted global correlation coefficient of 0.95 when compared against the reference diversity patterns of vascular plants. All values are averages across all botanical countries. Values are rounded up to three decimals. SE = standard error, CV = (ratio of sd to mean) *100

| **Sample** | **Diversity metric** | **Rank difference** | **CV (%)** | **SE** |
| --- | --- | --- | --- | --- |
| **1,048 spp.** | Species richness | 3.41 | 67.218 | 0.266 |
| **1,048 spp.** | Phylogenetic diversity | 11.16 | 57.937 | 31.498 |
| **1,048 spp.** | E. growth form diversity | 75.158 | 61.712 | 0.062 |
| **1,048 spp.** | E. family diversity | 70.326 | 66.473 | 0.199 |
| **67,772 spp.** | Species richness | 0.473 | 7.851 | 1.938 |
| **67,772 spp.** | Phylogenetic diversity | 4.946 | 7.104 | 80.046 |
| **67,772 spp.** | E. growth form diversity | 8.351 | 6.942 | 0.017 |
| **67,772 spp.** | E. family diversity | 23.788 | 9.194 | 0.229 |

### Figures

Figure SI1 Correlation of the coefficient of variation and the average diversity rank of a botanical country in 100 random samples extracted at two sample sizes (sample sizes correspond to the lowest and highest number of species needed to reach a geographically weighted global correlation coefficient of 0.95 when compared against the reference diversity patterns of vascular plants). Top right displays the spearman correlation coefficient for each sample size and the associated p-value.

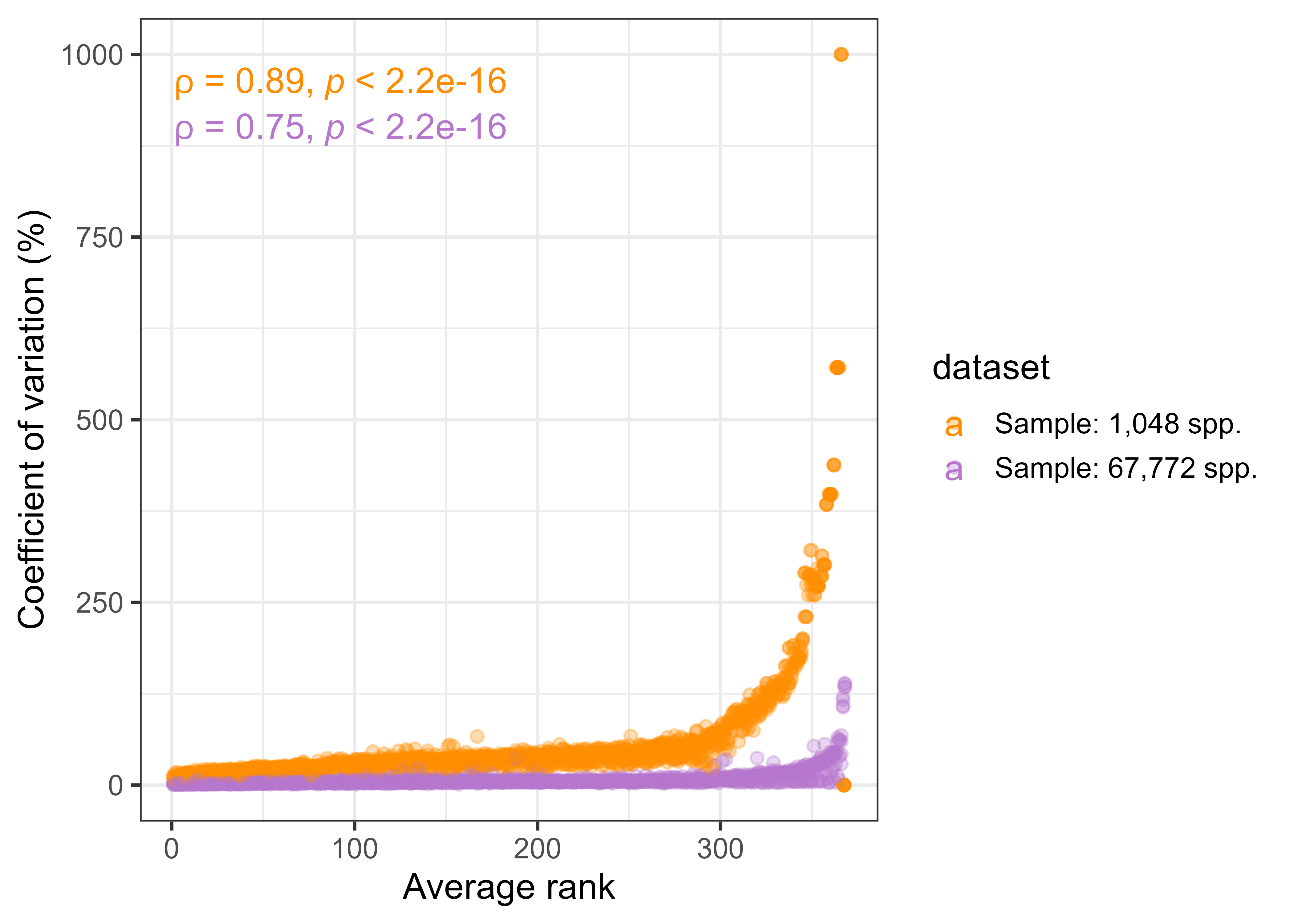
